# Weak evidence for change in wind-induced bending moments on raised and thinned Colorado spruce (*Picea pungens*)

**DOI:** 10.64898/2026.06.11.731663

**Authors:** David Leinbach, Daniel C. Burcham, Brian Kane

## Abstract

Trees are routinely pruned to mitigate the risk of wind damage, but there are few studies examining changes in wind loads after pruning, especially for large conifers. In this study, ten Colorado spruces (*Picea pungens*) were monitored before and after a series of pruning treatments. Trees were pruned to raise or thin crowns over a range of severities between 0% and 40%. Wind-induced bending moments were measured using two calibrated displacement probes installed orthogonally on the lower stem of each tree. Using a hierarchical Bayesian model, the relationship between maximum wind speeds and bending moments was quantified, consistent with theoretical and empirical expectations, as a non-linear power law. Random intercepts for model coefficients were used to account for individual variability in aerodynamic behavior among experimental trees, and predictions were made using the median response marginalized over the observed trees. The modeled relationship between wind speeds and bending moments was physically reasonable and like existing measurements with scaling exponents below two. Despite considerable variation among experimental trees, the aerodynamic behavior of trees, as indicated by model coefficients, was not clearly altered by pruning treatments, and, correspondingly, model predictions of bending moments over the range of observed wind speeds remained similar for all pruning treatments. Ultimately, the study yielded weak evidence for a change in bending moments following conventional pruning treatments for Colorado spruce, and the practical value of pruning to mitigate risk appeared limited for the studied conditions.

**Highlights:**

- Wind loads were monitored on large Colorado spruce after crown raising and thinning
- A hierarchical Bayesian model quantified wind speed and bending moment power laws
- Negligible change in bending moments was found for all pruning types and severities
- Conventional pruning methods may not mitigate risk for Colorado spruce

## Introduction

Often used to improve structure, enhance aesthetics, maintain clearance, or mitigate risk, pruning is one of the most common tree maintenance practices in communities around the world. However, there is limited research on the various outcomes of pruning treatments, especially changes in wind loads. Consequently, the industry guidelines developed for arborists have largely been adopted without thorough empirical testing (“ANSI A300 Tree Care Standards,” 2023). Lacking clear empirical evidence to guide them, arborists must subjectively remove branches to achieve management goals without harming the tree, but the most appropriate pruning type and severity are usually not clear for most objectives, except for simple tasks like maintaining clearance or removing deadwood.

In particular, risk mitigation is a common motivation for pruning large, mature trees. Although arborists often aim to reduce wind loads by pruning, few studies have examined the effects of pruning on the interaction between wind and trees (Gilman et al., 2008a, 2008b; Pavlis et al., 2008; Smiley and Kane, 2006). Moreover, methodological inconsistencies and the use of small, deciduous trees in existing studies have limited generalization about results to practical settings. One recent study measured ambient wind loads on large Senegal mahoganies (*Khaya senegalensis*) after pruning, and the authors reported that bending moments decreased more on reduced than raised trees over a range of pruning severities (Burcham et al., 2021). For the reduced trees, wind loads decreased quickly at relatively low pruning severities (≤ 20%) alongside favorable changes in vibration properties (Burcham et al., 2021, 2020). The findings were consistent with some work on smaller trees (Pavlis et al., 2008; Smiley and Kane, 2006), but there is a need for additional measurements of the pruning-induced changes in wind loads for large trees, especially other species with distinct crown characteristics.

Despite their prevalence in many urban forests, there have been exceptionally few pruning studies involving coniferous evergreens. Unlike broadleaf deciduous trees, their needle leaves persist during the winter months, often during stronger wind conditions (Jung et al., 2022), and deform less during flow (Rudnicki et al., 2004; Vollsinger et al., 2005). Colorado spruce (*Picea pungens*) is widely planted in many areas, and many practitioners have observed its propensity to fail during strong winds in the winter months, possibly due to its higher crown density, stiff needles, or shallow root systems. Given their consistently excurrent crown architecture, they are rarely reduced by arborists, and the practical options for pruning to mitigate risk may be limited for Colorado spruce and similar species. Therefore, the objective of this study was to evaluate the effects of common pruning types and severities on the wind-induced bending moments of Colorado blue spruce.

## Methods

### Site and trees

Ten mature Colorado spruce were selected for this study from a border planting at the Colorado State Forest Service (CSFS) Nursery in Fort Collins, CO, USA (Figure 1). The planting contained two parallel rows of trees oriented roughly north-south, including one row of 18 Colorado spruces on the western edge and another row of 22 ponderosa pines (*Pinus ponderosa*) on the eastern edge. The trees had not been previously pruned, and their form was typical of open grown spruce with an excurrent, conical crown extending from the ground to the top of the tree.

**Figure 1:**
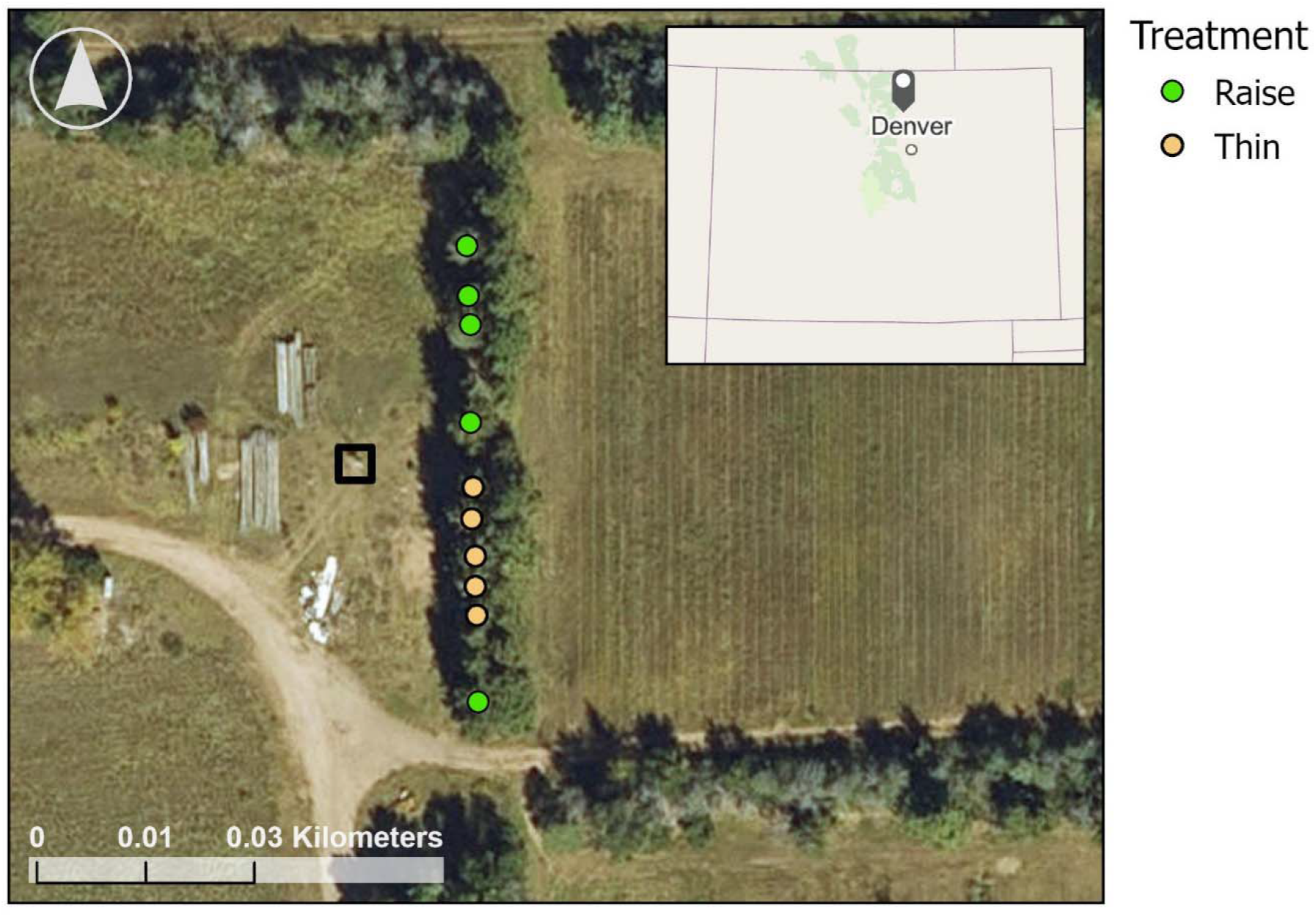
Ten Colorado spruce (*Picea pungens*) were selected for study from a windrow at the Colorado State Forest Service Nursery in Fort Collins, CO (gray marker, inset map). During the pruning experiment, the tree crowns (circle markers) were either raised (green) or thinned (orange). Wind conditions were monitored with an anemometer attached to a guyed mast (black square) located west of the study trees. Following a decline in its condition, the tree at the northern end of the row in the depicted layout was removed from the study.

Large, healthy trees with minimal dieback and even branch spacing were selected for the experiment. Subsequently, the neighboring trees not selected for the study were pruned to create additional clearance around the study trees and prevent collisions during wind events. Several morphological characteristics of each spruce were recorded, including tree height, crown length, and diameter at breast height (DBH) measured 1.37 m above ground (Table 1). The age of the trees was unknown because planting records did not exist.

**Table 1:** Morphological and mechanical measurements for 10 Colorado blue spruce (*Picea pungens*) used as experimental trees.

| Tree | Diameter (cm) | Height (m) | Crown length (m) | Stiffness |  |  |
| --- | --- | --- | --- | --- | --- | --- |
| | | | | Young's Modulus, $E$ (GPa) | Calibration coefficient, $C_1$ (kNm·mm<sup>-1</sup>) | Flexural rigidity, $EI$ (MN·m<sup>2</sup>) |
| 1 | 46.8 | 13.3 | 13.3 | 2.9 (0.02) | 83.0 (0.6) | 7.2 (0.05) |
| 2 | 36.9 | 12.0 | 11.8 | 2.3 (0.01) | 30.3 (0.1) | 2.0 (0.01) |
| 3 | 36.4 | 13.7 | 13.7 | 3.0 (0.05) | 41.9 (0.7) | 3.1 (0.05) |
| 4 | 39.5 | 15.2 | 15.2 | 4.8 (0.05) | 85.9 (0.9) | 6.9 (0.07) |
| 5 | 39.2 | 13.5 | 12.1 | 4.6 (0.07) | 75.0 (1.1) | 5.4 (0.08) |
| 6 | 37.7 | 12.7 | 11.9 | 4.5 (0.04) | 65.3 (0.6) | 4.6 (0.04) |
| 7 | 43.4 | 13.7 | 13.7 | 4.2 (0.03) | 92.8 (0.6) | 7.2 (0.05) |
| 8 | 34.6 | 10.8 | 10.6 | 2.2 (0.02) | 25.7 (0.2) | 1.7 (0.01) |
| 9 | 45.3 | 13.7 | 13.7 | 4.3 (0.02) | 105.7 (0.5) | 8.8 (0.05) |
| 10 | 45.9 | 11.9 | 11.8 | - | - | - |
| Mean | 42.5 (4.4) | 13.1 (1.2) | 12.8 (1.4) | 3.6 (1.0) | 67.3 (28.4) | 5.0 (2.5) |

Note: Stiffness values for individual trees are mean (SD) for three separate trials; bottom row contains mean (SD) computed for all trees; tree 10 was removed from the experiment after developing extensive foliar necrosis.

### Instrumentation

A three-dimensional ultrasonic anemometer (Model 81000, R.M. Young Company, Traverse City, MI, USA) was used to measure wind velocity. The anemometer was installed on a 14.6 m weather mast (PA2 – Transportable Mast System 3-Way Halyard, South Midlands Communications Ltd., Eastleigh, Hampshire, England) at the average height of experimental trees (13.1 m). The weather mast was erected 15 m west from the middle of the row of experimental trees (Figure 1), and the anemometer was mounted on a 1 m horizontal boom oriented northeast to minimize flow distortion during prevailing northwestern winds. A lightning protection system was installed on the mast to mitigate the risk of equipment damage.

Axial trunk displacement, *d*(mm), was monitored by two linear variable displacement transducers (LVDT) (VS/20/UU/5, Solartron Metrology, West Sussex, UK) mounted orthogonally on the stems at 1.37 m above grade. Before use, the strain resolution of each LVDT was approximately doubled from 44 μm·m^−1^ to 23 μm·m^−1^ by adding a 200 mm segment of threaded rod to the LVDT body. To minimize solar heating, the LVDTs were mounted on the east and north sides of each stem. Secured to the trees using hanger bolts, the probes measured both elongation and contraction of the outer wood during wind events.

Wind velocity, ***u***(m·s^-1^), and *d* were simultaneously and continuously recorded at 20 Hz from July 2023 to January 2024 at the experimental site. The 20 LVDT probes and anemometer were wired to analogue and digital channels, respectively, on two dataloggers (CR1000 and CR1000X, Campbell Scientific Inc., Logan, UT, USA) mounted on the weather mast inside a weather resistant enclosure (ENC10/12, Campbell Scientific Inc., Logan, UT, USA). The entire system was powered by a sealed lead acid battery (40701, Grainger, Kansas City, MO, USA) housed in an insulated battery box (HM-484 by The NOCO Company, Glenwillow, OH, USA) to protect against temperature fluctuations. Battery charge was maintained through a 90-watt solar panel (SP90-L15-P, Campbell Scientific Inc., Logan, UT, USA) mounted 0.5 m above the weather resistant enclosure on the mast, and power management was controlled through a charge regulator (CH200, Campbell Scientific Inc., Logan, UT, USA).

To estimate wind-induced bending moments, *M*_*B*_ (kN·m), from measured displacements, a calibration coefficient, *C*_*1*_ (kN·m·mm^-1^), was determined for the pair of LVDTs mounted on each experimental tree using a series of static pull tests (Wellpott, 2008). To increase the applied force and slow the rate of loading, the trees were pulled using a 2:1 rigging system aligned incident to the eastern LVDT on each tree. A rope (9/16 in Stable Braid Bull Rope, Samson Rope Technologies, Ferndale, WA, USA) was attached to a truck-mounted winch (VR Evo 8-S, Warn Industries, Inc., Clackamas, OR, USA), routed around the sheave of a block (10 Ton Snatch Block, TICONN, China) attached to the tree, and terminated under the winch on the truck. A load shackle (ISHKB3.25MT1361, Interface Force Measurement Solutions, Scottsdale, AZ, USA), connecting the block to a dead-eye sling (5/8 in Stable Braid Bull Rope, Samson Rope Technologies, Ferndale, WA, USA) attached to the tree stem, measured the total tensile force generated by the rigging system during testing. During testing, loads were recorded on a synchronized analogue channel in the data acquisition system used for other measurements. The incremental *M*_*B*_ generated at the middle of the displacement probe was calculated as:

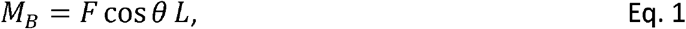

where *F* is the force (kN) applied by pull rope, *θ* is the angle between the rope and a horizontal plane, and *L* is the vertical distance (m) between the pull rope attachment point and the middle of the displacement probe. *C*_1_ was estimated as the slope of an ordinary least-squares regression line fit to model *M*_*B*_ as a function of *d*.

To estimate trunk stiffness, Young’s modulus, *E* (N·m^-2^ = Pa), was calculated as:

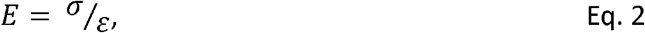

where σ (N·m^-2^ = Pa) is stress, calculated as:

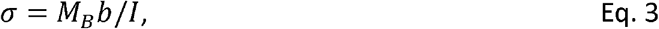

where *b* is the trunk radius parallel to the direction of bending and *I* is the second moment of area (m^4^) determined using:

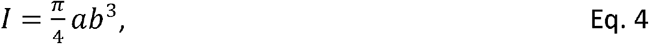

where *a* is the trunk radius perpendicular to the direction of bending. Strain, *ε* (dimensionless), was calculated as:

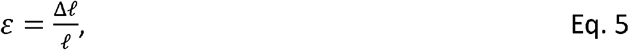

where Δ*ℓ* is the change in length of the displacement probe and *ℓ* is the undeformed length of the LVDT body.

### Pruning treatments

Trees were either raised or thinned broadly according to industry standards over a range of severities. Trees were randomly assigned to a pruning type (i.e., raised, thinned), and five different severities were progressively applied to the same individual trees during the experiment, including 0%, deadwood removal, 10%, 20%, and 40%. The removal of dead branches was not quantified, but all trees contained a similar and substantial amount of deadwood.

On raised trees, branches were removed to increase vertical space between the crown and ground. The severity of pruning was calculated as the percent change in crown height, *H*_*CROWN*_ (m):

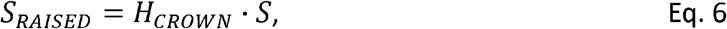

where *H*_*CROWN*_ is the vertical distance (m) between the crown apex and lowest branch and *S* is the target severity (%).

On thinned trees, branches were selectively removed to increase the crown porosity, and the severity of pruning was determined using the cumulative sum of branch diameters, Σ_*DIA*_ (m), broadly according to Kane et al. (2014). All primary branch diameters were recorded for each tree using a dial caliper (D15TX, Mitutoyo Corporation, Takatsu-ku, Kawasaki-shi, Kanagawa, Japan). Starting from the lowest branch, measurements were recorded until the trunk diameter decreased below 9.5 cm. Total severity was calculated as:

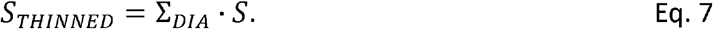

The initial crown density was not uniform due to branch and needle senescence in older branches near the ground. To create a more uniform crown density profile, half of the target severity was applied to the top third of the crown, and the remaining half was split between the bottom two thirds.

### Data processing and statistical analysis

30-minute data intervals were consistently used for analysis to capture typical variation in wind conditions while avoiding problematic noise and fluctuations at shorter time intervals and slowly varying trends at longer time intervals. First, the quality of time histories was checked using various signal time and frequency features. Trunk displacement outliers were identified as values more than three scaled median absolute deviations from the median value within a sliding window of 250 values, and they were replaced, as needed, with the nearest non-outlier value. Wind speed outliers were similarly detected and replaced. To ensure that measured displacements did not erroneously exceed the physical measurement range of each probe, intervals were removed if the measured range exceeded 20 mm. To compare the actual with expected measurement resolution of probes, time histories were numerically differentiated to confirm agreement with expectations for the sensor and data acquisition system, and intervals with incorrect measurement resolution were removed.

To detect and remove trunk displacement time histories contaminated by noise, an ensemble average power spectral density (PSD) was computed using eight separate non-overlapping windows with 4,096 observations multiplied by a Hamming window to minimize spectral leakage. Since trees vibrate in the decihertz range (de Langre, 2019), the absence of a prominent peak in PSD between 0.1 and 1 Hz for time intervals with an average wind speed above 1.5 m·s ^-1^ was assumed to be caused by noise, and the time history was excluded from further analysis.

Based on the assumption that tree failure is an extreme value process, the maximum of several statistics was computed for each retained 30-minute interval for modeling. For 30-minute intervals with *n* observations, the maximum resultant bending moment, *M*_*B,max*_ (kN·m), was calculated as:

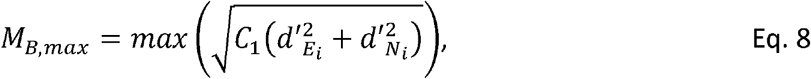

where 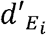 and 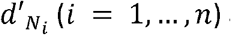 are the stem displacements measured along the east-west and north-south directions, respectively, and fluctuating about their respective mean values for each tree. Maximum horizontal wind speed, *U*_*max*_ (m·s^-1^), was computed as:

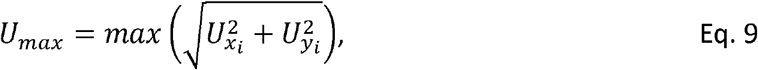

where 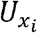 and 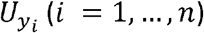 are the instantaneous wind speeds measured along the anemometer’s horizontal *x* - and *y*-axes, respectively. Maximum friction velocity, *u* *_*max*_ (m·s^-1^), was computed as:

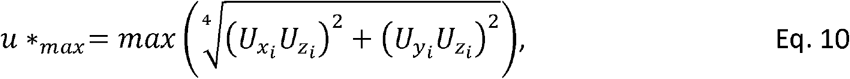

where 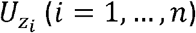 are the instantaneous wind speeds measured along the anemometer’s vertical *z*-axis.

Consistent with theoretical expectations (Angelou et al., 2019), the relationship between *M*_*B,max*_ and *U*_*max*_ was modeled as a non-linear power function. A lognormal likelihood with an identity link was used to account for positively skewed, heteroscedastic *M*_*B,max*_. The model was defined as:

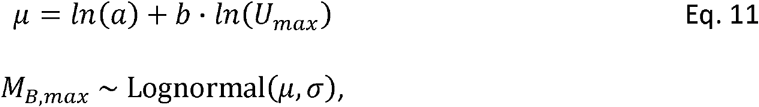

where *a* and *b* are the scaling coefficient and exponent, respectively, for the power function, and *μ* and *σ* are the linear predictor and standard deviation on the log scale, respectively. Pruning treatments were modeled as fixed effects of both *a* and *b*. A random intercept for *a* and *b* was used to account for variability among individual trees.

The independent variable *U*_*max*_ was selected through a formal comparison of candidate models against *u* _* *max*_. To determine the fundamental coupling between wind conditions and tree response, the models were fit to data, unconfounded by pruning treatments, collected during the 0% severity period, and the candidates were compared using Bayesian Leave-One-Out Cross-Validation (LOO-CV) to compute the expected log-predictive density (ELPD) for each model. The *U*_*max*_ model showed the highest posterior predictive accuracy (ΔELPD > 170), and it was used consistently in all analyses.

Using Bayesian inference, the model was fit in Stan (Carpenter et al., 2017) using the brms package (Burkner, 2017) in R version 4.2.1 (R Core Team, 2025). To avoid excessively constraining parameter estimates, weakly informative priors were set using informed theoretical expectations and prior predictive checks: *a* ~ Normal(2.0.5,1), *b* ~ Normal(2.0,0.5), *σ* ~ Exponential(1), and an LKJ(2) prior for the correlation between random effects. Posterior distributions were sampled using four independent MCMC chains with 2,000 warmup and 2,000 post-warmup iterations each. The target acceptance rate and maximum tree depth were set to 0.999 and 15, respectively, to promote robust sampling and chain convergence.

Model validation was performed using posterior predictive checks to confirm that the simulated distribution of *M*_*B,max*_ accurately reflected the observed data. The influence of individual observations on posterior estimates was evaluated using the Pareto *k* statistic from LOO-CV on the full model, and observations with *k* > 7 were considered high-leverage and carefully examined for their impact on model stability. The model’s predictive precision was assessed using the mean absolute error, MAE (kN·m), between observed and predicted *M*_*B,max*_ from all post-warmup posterior draws. During inference, treatment predictions were marginalized over the tree-level random effects to represent the median loading response of a typical individual from the studied group of trees.

## Results

During the initial phase of data collection, one tree was removed from the study due to extensive foliar necrosis in the lower portion of the crown. As a result, the number of experimental trees in the raised treatment group was reduced to four, and all other trees remained healthy and persisted throughout the remainder of the experiment. For the remaining nine trees, *E* determined from load tests varied over a 2.4 GPa range between 2.2 and 4.8 GPa (Table 1), and C_1_ spanned the low multiples of 10 kN·m·mm^− 1^. Estimates derived from multiple load tests were very consistent with low variability among replicate measurements (Figure 2).

**Figure 2:**
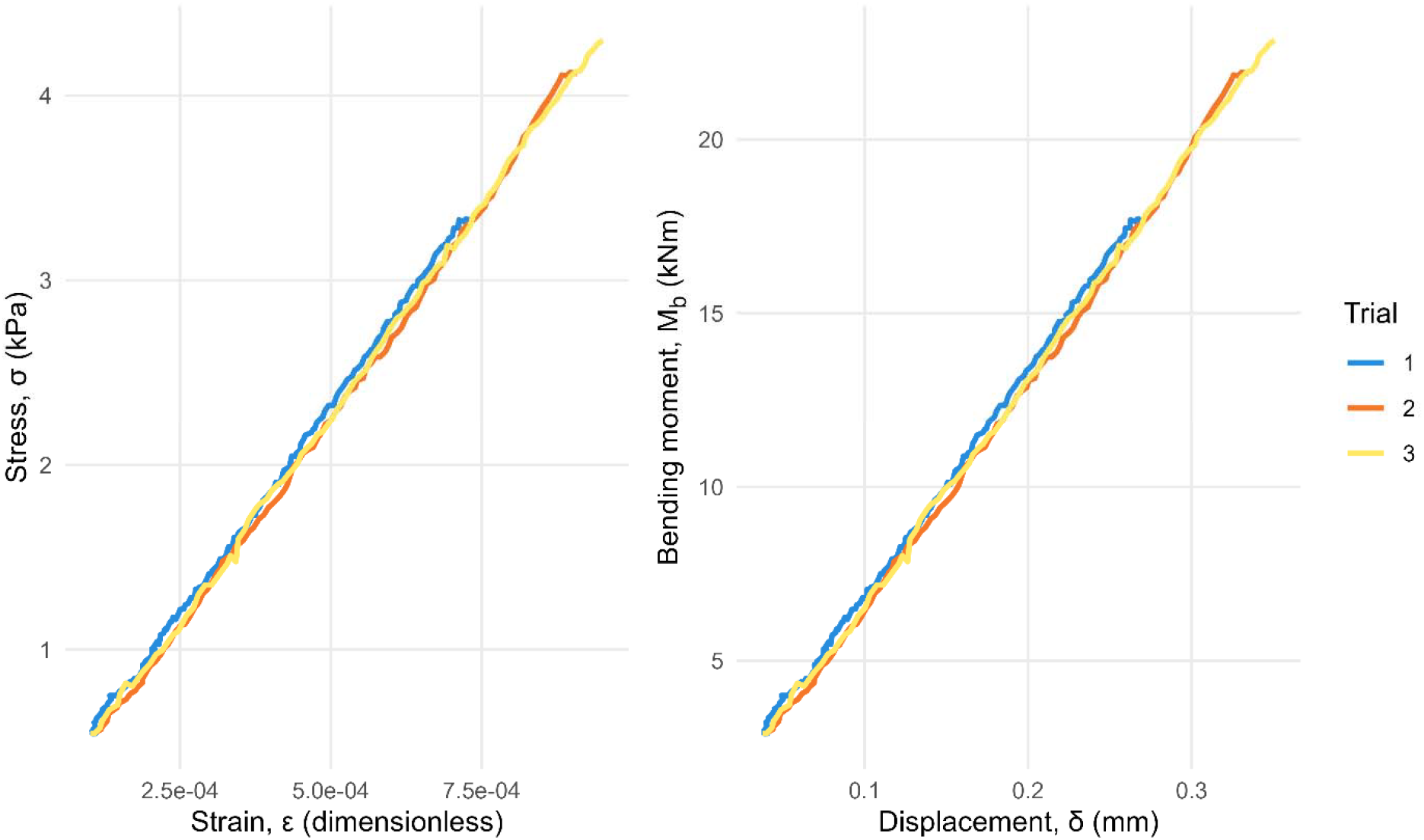
Static pull tests were used to determine the cross-sectional stiffness of each Colorado spruce (*Picea pungens*) stem 1.37 m above ground with stress-strain (left) and momentdisplacement (right) plots. Using illustrative data from tree #6, the slope of a line fit to data from each test trial was used to estimate Young’s modulus, *E* (GPa), and calibrate sensors for measurements, *C*_1_(kNm·mm-1). For estimates from all trees, see Table 1

At the start of the experiment, data were collected for an extended 50-day period, but the remaining pruning treatments were subsequently delayed by a malfunctioning charge regulator in the data acquisition system. After repairs, the experimental schedule resumed, but the schedule was accelerated to minimize data collection during freezing winter temperatures. For the altered schedule, the treatments were maintained for at least one week to characterize tree responses to a range of wind conditions. The deadwood removal, 10%, 20% and 40% pruning severity treatments were monitored for 10, 12, 8, and 28 days, respectively. After excluding signals contaminated by noise, 15,640 30-minute intervals from all experimental periods were used for analysis. In total, 8,486, 1,759, 926, 1,039, and 3,430 30-minute intervals were used from the 0%, deadwood, 10%, 20%, and 40% severities, respectively.

Wind conditions were relatively calm during the experiment. Ensemble mean (range) 30-minute mean and maximum horizontal wind speeds were 2.0 m·s^-1^ (0.3 – 7.7) and 5.3 m·s^-1^ (0.3 – 16.8), respectively. Turbulent fluctuations were similarly moderate; ensemble mean (range) 30-minute mean and maximum friction velocities were 0.5 m·s^-1^ (0 – 1.8) and 2.3 m·s^-1^ (0 – 8.8), respectively. Wind conditions were broadly similar during sequential experimental phases (Figure 3). During each phase, the prevailing (modal) winds flowed towards easterly directions, and 30-minute maximum wind speeds spanned a similar range between roughly 1 and 16 m·s ^-1^.

**Figure 3:**
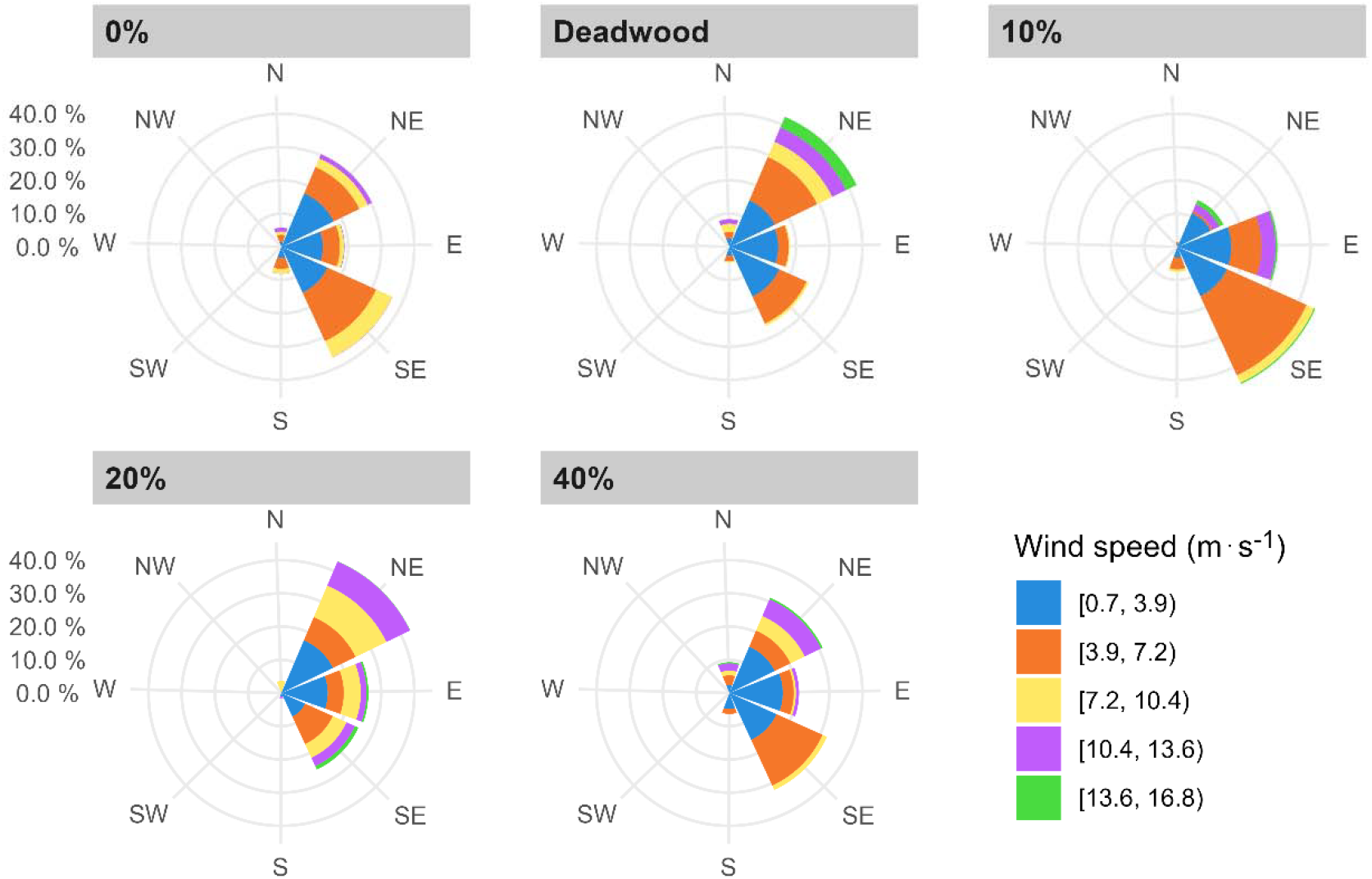
Wind roses showing 30-minute maximum horizontal wind speeds, *U*_*max*_ (m·s-1), and prevailing (modal) directions. For each experimental period, the length of spokes shows the relative frequency of wind directions, and the thickness of colored bands shows the distribution of speeds at each direction.

Visual inspection of trace plots indicated good sampling of the parameter space by MCMC chains. All parameters reached convergence with 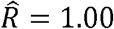, and sampling efficiency was high with bulk effective sample size (ESS) ratios exceeding 0.24 for all parameters. Posterior predictive checks showed close agreement between the modeled and empirical distribution of *M*_*B,max*_ (Figure 4). The model exhibited high predictive precision with a median MAE of 0.76 kN·m (95% PPI: 0.75 – 0.77 kN·m). Furthermore, all observations exhibited k < < 0.7, with the majority below 0.4, confirming an insensitivity to outliers. Consistent with stiffness measurements (Figure 5), model estimates for individual trees conformed to their underlying data, with a lower *M*_*B,max*_ at a given *U*_*max*_ for relatively flexible trees.

**Figure 4:**
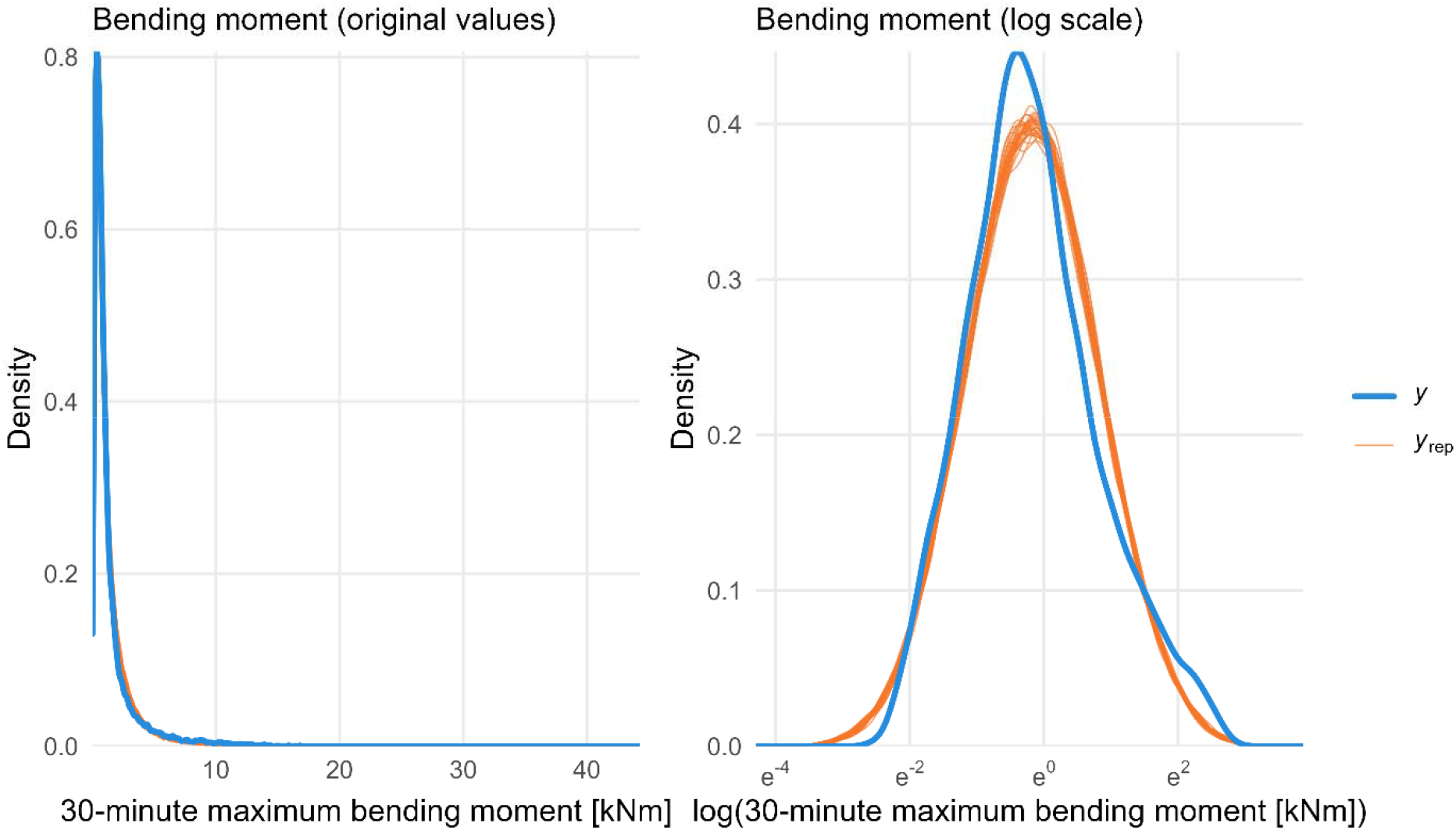
Posterior predictive check showing good agreement between 30 posterior simulated (orange) and observed (blue), *M*_*B,max*_ distributions in the linear (left) and logarithmic (right) scales

**Figure 5:**
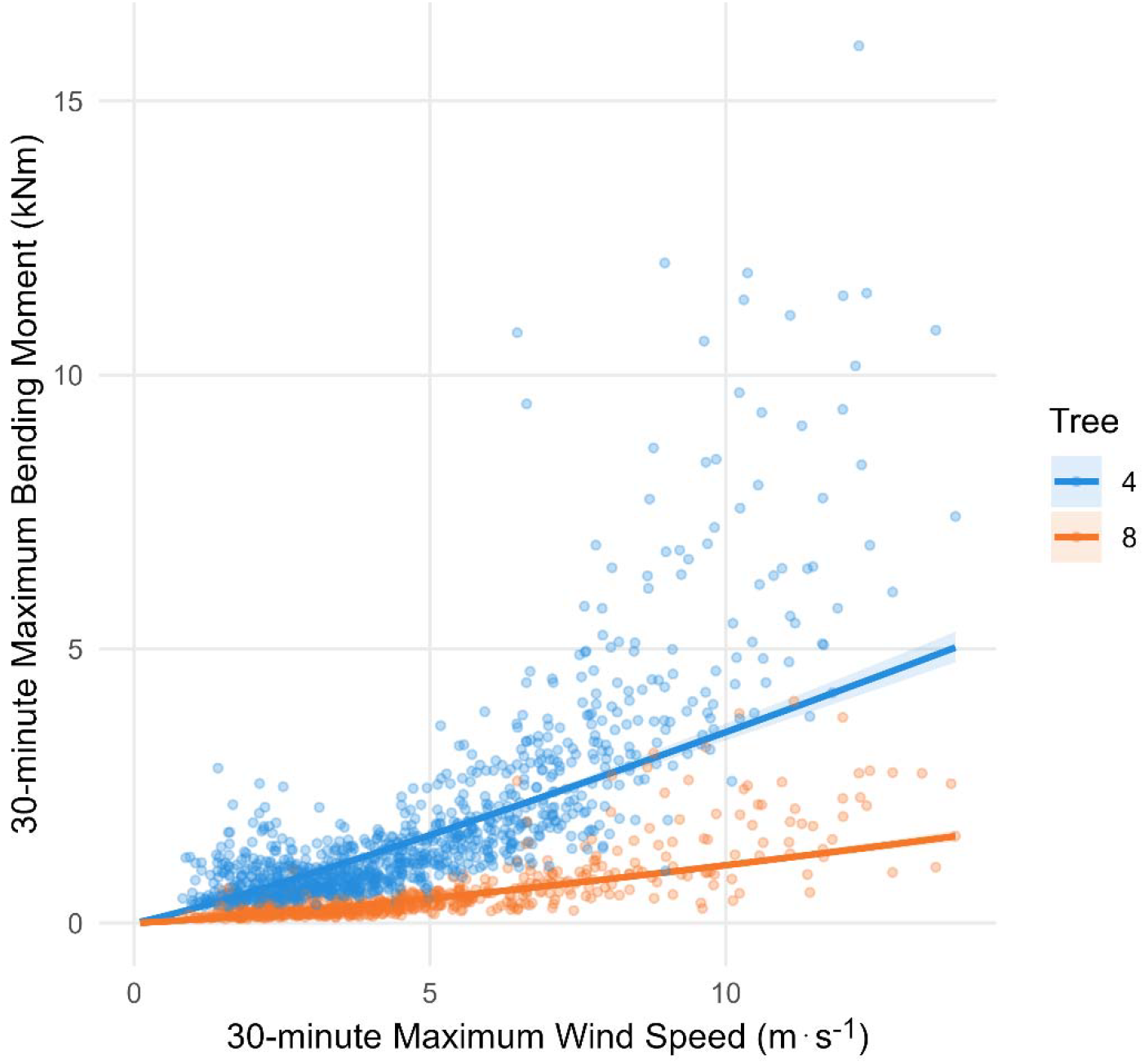
Predicted relationship between 30-minute maximum bending moment, *M*_*B,max*_ (kNm), and 30-minute maximum wind speed, *U*_*max*_ (m·s-1) for two experimental trees with the highest (Tree 4: 4.8 GPa) and lowest (Tree 8: 2.2 GPa) cross-sectional stiffness measured during static load tests. Solid lines represent conditional posterior predictive medians, and shaded regions indicate 95% credible intervals

Model coefficients remained substantially similar across pruning treatments with scaling factors near 1.2 and exponents between 1.5 and 1.8 (Table 2), and the 95% credible intervals for the parameters covered an equivalent, broad range of values for all pruning treatments. The model also revealed a weakly negative correlation (*r* = −0.21) between the tree-level random effects for the scaling coefficient (*a*) and exponent (*b*). The continuous relationship between *M*_*B,max*_ and *U*_*max*_, derived from the marginal posterior predictive distribution of *M*_*B,max*_, represented the median loading profile of experimental trees for each pruning treatment (Figure 6). While the 0% severity curves generally showed lower bending moments at a given wind speed compared to other treatments, the 95% prediction intervals for all pruning treatments, incorporating both parameter uncertainty and residual error, were largely coincident throughout the observed wind speed range. Near the upper limit of observed wind speeds (*U*_*max*_ = 15 m·s^-1^), the absolute difference in marginal posterior median *M*_*B,max*_ between 0% and all other severities was less than 5 kN·m for both pruning types (Figure 7), and the 66% prediction intervals for the differences mostly encompassed zero, indicating that the treatment effect was small compared to total predictive uncertainty.

**Table 2:** Model coefficients for power laws fit to 30-minute maximum bending moment (*M*_*B,max*_) and 30-minute maximum horizontal wind speed (*U*_*max*_) for all pruning treatments.

| Pruning Severity | Model Coefficients ($M_{B,max} = aU_{max}^b$) | |
| --- | --- | --- |
| | Scaling Factor ($a$) | Scaling Exponent ($b$) |
| **Raised** |  |  |
| 0% | 1.19 [1.08, 1.32] | 1.63 [1.24, 2.02] |
| Deadwood | 1.19 [1.07, 1.32] | 1.73 [1.34, 2.12] |
| 10% | 1.19 [1.07, 1.32] | 1.73 [1.33, 2.12] |
| 20% | 1.19 [1.07, 1.32] | 1.80 [1.40, 2.19] |
| 40% | 1.19 [1.07, 1.32] | 1.78 [1.39, 2.17] |
| **Thinned** |  |  |
| 0% | 1.20 [1.09, 1.31] | 1.59 [1.16, 1.99] |
| Deadwood | 1.18 [1.08, 1.29] | 1.71 [1.29, 2.12] |
| 10% | 1.21 [1.11, 1.32] | 1.53 [1.11, 1.94] |
| 20% | 1.17 [1.07, 1.27] | 1.83 [1.41, 2.24] |
| 40% | 1.18 [1.08, 1.29] | 1.67 [1.24, 2.07] |

Note: Values are posterior medians and 95% credible (closed) intervals

**Figure 6:**
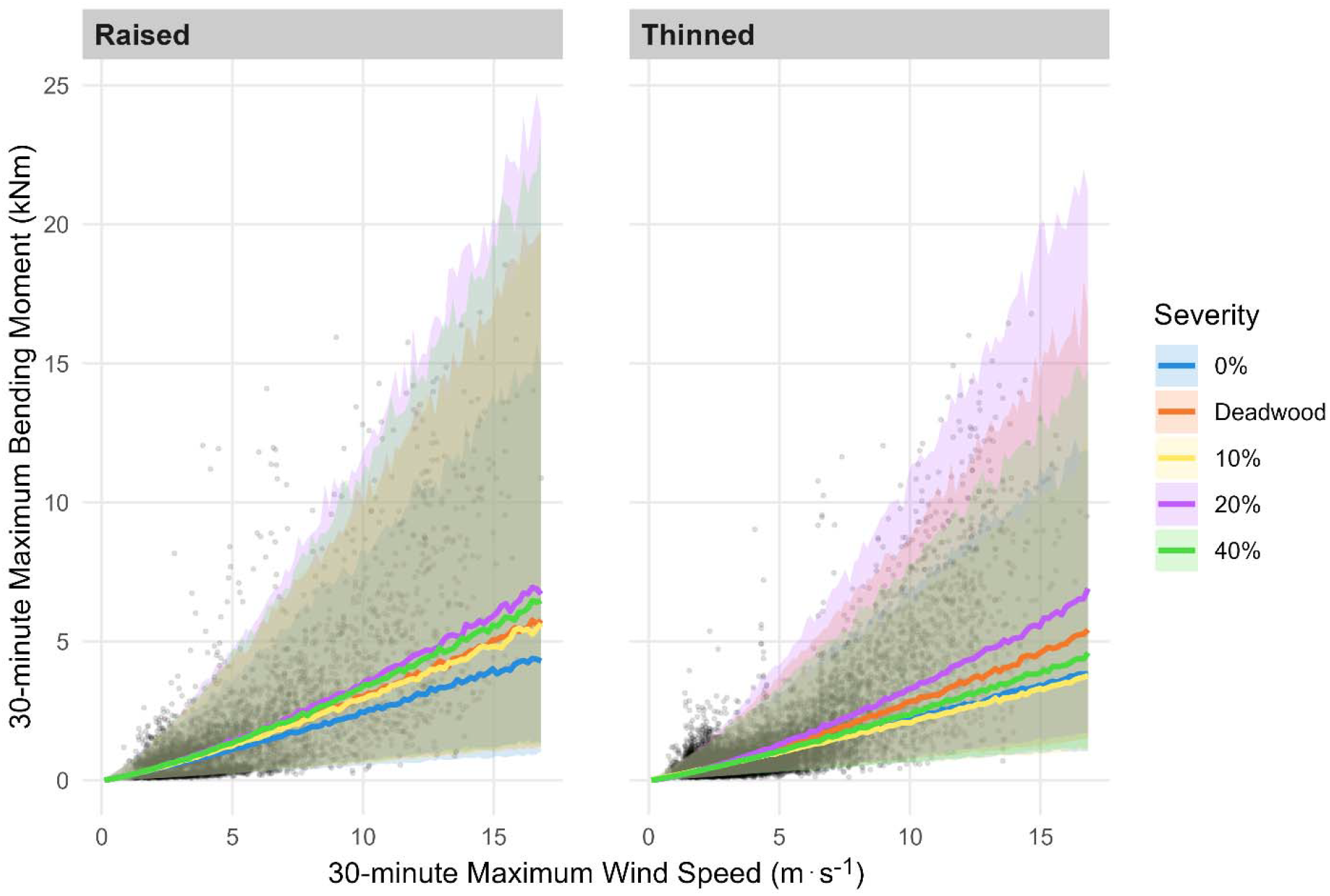
Predicted relationship between 30-minute maximum bending moment, *M*_*B,max*_ (kN·m), and 30-minute maximum wind speed, *U*_*max*_ (m·s-1) for two pruning types and five severities. Computed using posterior draws for all experimental trees, solid lines represent marginal posterior predictive medians, and shaded regions indicate 95% prediction intervals. Predictions were obtained from a hierarchical nonlinear Bayesian model fit to data from 15,640 30-minute intervals (black circle markers).

**Figure 7:**
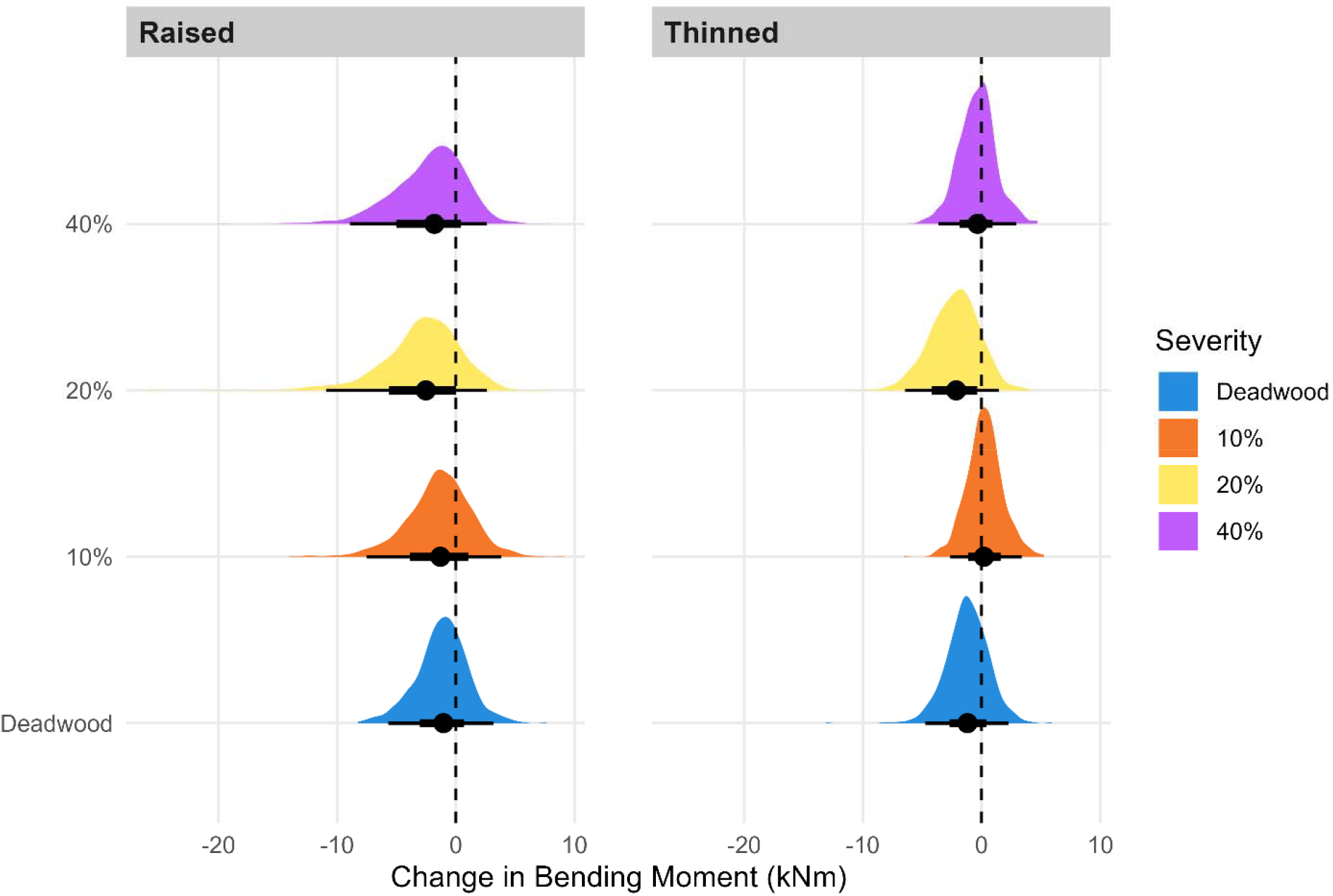
Marginal posterior predictive distributions of the difference in 30-minute maximum bending moment, *M*_*B,max*_ (kN·m), between the unpruned (0% severity) and four pruning treatments at *U*_*max*_ = 15 m·s-1. For raised (left) and thinned (right) trees, circle markers represent the posterior medians of the difference, and thick and thin horizontal bars indicate the 66% and 95% prediction intervals, respectively. Positive (negative) values indicate a decrease (increase) in, *M*_*B,max*_ and the vertical dashed line at zero indicates no change from the typical individual tree at 0% severit

## Discussion

Generally, the experiment yielded weak evidence that two conventional pruning types applied at common severities altered the aerodynamic behavior or peak mechanical loads of large Colorado spruce. The overlapping 95% posterior predictive intervals suggested that variability from individual trees, wind conditions, and measurement error outweighed the management intervention. Since wind-induced bending moments did not obviously decrease after pruning, the treatments studied may not be an effective risk mitigation strategy for Colorado spruce. However, there was also no evidence of a marked increase in bending moments on raised trees, despite speculation about such outcomes among practitioners, and the frequent removal of lower branches on the species for practical reasons does not likely cause adverse mechanical consequences.

While the scaling exponents were slightly higher than reported for smaller coniferous (Mayhead, 1973; Vogel, 1989) and broadleaf (Kane and Smiley, 2006) trees, they were similar to some measurements of large broadleaf trees during seasonal periods with leaves (Angelou et al., 2025, 2019). Equivalent to Vogel exponents, 𝒱, between −0.4 and −0.2, the scaling exponent values (*b* < 2) suggest that reconfiguration occurred, but it was less than observed for smaller, more flexible trees. The wood stiffness measured during load tests in this study were similar to reported measurements of other trees (Senalik and Farber, 2021), but the (unmeasured) stiffness of branches and leaves may have impeded greater reorientation during flow, possibly due to age or species characteristics. Relatively few studies have characterized variation in stiffness among different tree parts, but existing measurements show that patterns of variation in material properties strongly differ among species (Gurau, 2008; Woodcock and Shier, 2002). Despite considerable alterations from branch removal, the aerodynamic behavior of the trees apparently remained broadly similar after pruning.

Most existing studies have reported that bending moments decreased the most after reducing the height or spread of crowns during pruning (Burcham et al., 2021; Gilman et al., 2008a; Pavlis et al., 2008; Smiley and Kane, 2006). Since bending moments are the product of drag and center of pressure, the removal of distal branches in the upper crown simultaneously decreased both governing factors precipitously, but they were not similarly altered on raised or thinned trees. Given low crown density and wind speeds near the ground, the removal of lower branches likely caused a modest decrease in drag and slight increase in the center of pressure on raised trees, and the two changes may have had counteracting effects on bending moments (Pavlis et al., 2008). On thinned trees, the removal of whole branches reduced the density of crowns without altering total area, and the results indicated that drag remained similar after pruning. After pruning, the retained branches on thinned trees often had a sizeable proportion of dead needles and twigs, given the species low propensity for natural shedding, and the residual crown density may have contributed to persistent drag.

The species’ leaf and crown traits may have conferred a higher degree of mechanical tolerance, instead of aerodynamic avoidance, to wind loads (Puijalon et al., 2011). In this study, there was no evidence of abnormal stiffness in the lower stem compared to reported values for other species (Lundstrom et al., 2008; Senalik and Farber, 2021), but the lack of a pronounced change in the aerodynamic response of Colorado spruce could be partially explained by crown characteristics. In future studies, it will be important to carefully measure biomechanical traits (e.g., material properties) and track changes in crown characteristics (e.g., leaf area density, total area) during pruning to support analysis.

This study was constrained by several limitations that may have affected the results. The prevalent noise in analogue sensors excluded a large portion of data from analysis, and the improvement of instrumentation for noise reduction should be a main priority in future work. The use of grounded cable shielding, differential voltage measurements, and temperature correction should be adopted to minimize background noise. In addition, wind speeds were relatively calm during the entire study period with a maximum instantaneous horizontal wind speed of 16 m·s^-1^, and the relatively calm site conditions precluded an analysis of tree response to severe wind conditions. While a different experimental design maintaining all pruning treatments simultaneously on a larger group of trees for longer observation periods could increase the range of observed wind speeds, it could also be confounded by growth dynamics over time, especially if monitored for several months, and limited by higher instrumentation costs. The design of similar experiments deserves more careful consideration of such challenges in future work.

## Conclusion

For conventional pruning treatments, the change in wind-induced bending moments was modest for large Colorado spruce under mild wind conditions. Practically, the limited mechanical benefit of removing biomass may not justify the effort if load reduction is the primary objective. However, the study did not find evidence of an increase in wind loads after pruning, and the methods can likely be used for other practical reasons, such as maintaining clearance or managing wildfire fuels, without adverse mechanical consequences for the trees. Given their attractiveness and adaptability, conifers, such as Colorado spruce, are a substantial component of many urban forests. While practitioners generally avoid reduction pruning to preserve the excurrent architecture of conifers, there is ample evidence that crown reduction decreases wind loads on several broadleaf species, and the development of sensible approaches for reducing the size of conifers may be worthwhile when risk mitigation is a primary concern. Considering existing work, the study affirms that the mechanical effects of pruning differ among tree species with unique traits.

## Acknowledgments

The authors gratefully acknowledge Tony Koski and Yanlin Guo for feedback on an earlier version of this manuscript, Stuart Reinhoff and Sarah Wilhelm for support with field work, Jeff Smith and Emily Parsons for donating equipment, and the Colorado State Forest Service for permission to use the site and trees.

## CRediT Author Contribution Statement

**David Leinbach:** Conceptualization, Methodology, Investigation, Data Curation, Formal Analysis, Writing – Review & Editing. **Daniel C. Burcham:** Conceptualization, Methodology, Resources, Project Administration, Supervision, Software, Formal Analysis, Writing – Original Draft. **Brian Kane:** Supervision, Writing – Review & Editing.

## Data Availability

The data used in this study was deposited in the Harvard Dataverse at https://doi.org/10.7910/DVN/B0KQFU.

